# Dose assessment in dental cone beam computed tomography: comparison of optically stimulated luminescence dosimetry with the Monte Carlo method

**DOI:** 10.1101/675058

**Authors:** Chena Lee, Jeongmin Yoon, Sang-Sun Han, Ji Yeon Na, Jeong-Hee Lee, Young Hyun Kim, Jae Joon Hwang

## Abstract

The usage and the model variety of CBCT machine has been rapidly increasing, the dose evaluation of individual devices became an important issue. Patient dose from CBCT was assessed with two different methods, optically stimulated luminescence dosimeter (OSLD) measured and monte carlo (MC) simulation, in four different examination modes. Through the measurement process and obtained value, more practical and efficient method in acquiring CBCT effective dose would be suggested. Twenty-five OSLD were calibrated and equipped in human phantom of head and neck organs. This was exposed on 2 CBCT units, CS9300 (Carestream Dental LLC, Atlanta, Georgia) and RAYSCAN α+ (Ray Co. Ltd, Hwaseong-si, Korea) with 2 different examination modes. Dose recorded in dosimetry was obtained and organ dose as well as an effective dose were obtained in each units of examination modes. Those values were also calculated using MC software, PCXMC (STUK, Helsinki, Finland). The organ doses and effective doses from both methods were compared by each examination mode of individual unit. OSLD measured effective dose value was higher than that obtained with MC method in each examination mode, except dual jaw mode of CS9300. The percent difference of effective dose between the two methods were ranged from 4.0 to 14.3 %. The dose difference between the methods was decreased as the examination FOV decreased. Organ dose values were varied according to the method, while overall trend was similar in both methods. The organs showing high dose were mostly consistent in both methods. In this study, the effective dose obtained by OSLD measurement and MC simulation were compared and both methods were described in detail. Consequently, as relatively efficient and easy-handling method, we carefully suggest MC simulation for further dose evaluation.

## Introduction

Radiation in dental diagnostic examination is relatively low compared to that of medical [1, 2]. However, as cone beam computed tomography (CBCT) became largely performed for various purposes in dental clinics, we cannot say radiation dose in dentistry is very low any more.

Although patient’s overall radiation dose increased in dentistry, dose evaluation method is not developed very much. Dose measurement with thermoluminecent dosimetry (TLD) has been traditional method and most dental radiation dose researches were based on this method up to date [3]. There is recent trend of displacing TLD with optically stimulated luminescence dosimeter (OSLD) or metal oxide semiconductor field effect transistor (MOSFET) [4, 5]. MOSFET provides fast reading of dosage as it connected to the electronic probe directly. While it has been widely acceptable for dosimetry in radiotherapy, due to its suitability for high range of dose [6].

The basic phenomenological fundamentals of OSLD and TLD are the same while the TLD releases the energy, which was stored during irradiation, by heat and OSLD dose by light [7]. There are several advantages of OSLD over TLD such as, high sensitivity, preciseness and simple dosimeter preparation and readout. Based on these, a few literatures performed dose measurement with OSLD and they reported it showed reliable result compared to the TLD method [8]. Still, TLD has been a common dosimetry in dental field for a long time and there are not many studied based on OSLD measurement, yet.

Monte carlo (MC) method is another dose assessment method which simulate x-ray photon interaction with body organs and calculate overall effective dose. This method simulates virtual photon interaction on human phantom and expect radiation dose. Such method is advantageous in that it is simple to use since calibration and readout procedure are not required and the result is not dependent on the dosimeter types or its location in phantom [9]. However, this simulation is correct when it is based on the correct machine and radiation beam geometry, such as distance between the x-ray source to patient, beam rotation angle or vertical angle of x-ray beam. According to the incorrect combination of those factors, effective dose might show up to 51.24% difference compared to the TLD measured value [10].

Both OSLD measurement and MC simulation method are the short of data reported in dental x-ray equipment at present [10. 11]. More research on the newly introduced method, OSLD or MC calculation, compared to the traditional dosimetry should be performed to prove efficiency of these methods. In fact, as far as the authors know, there are no English reported study on dose assessment in comparison of MC method and OSLD method in the dental field.

In this study, patient dose from CBCT was assessed with two different methods, OSLD measured and MC simulation, in two different CBCT units with different examination modes. Through the measurement process and obtained value, more practical and efficient method in acquiring CBCT effective dose would be suggested.

## Material and methods

### 1. Cone beam CT (CBCT) machines and examination protocols

The CBCT equipment used were CS9300 (Carestream Dental LLC, Atlanta, Georgia) and RAYSCAN α+ CBCT (Ray Co. Ltd, Hwaseong-si, Korea).

The examination modes of individual units used in this study were as followed; In CS9300, facial mode (FOV = 17x 13.5 cm), dual jaw mode (FOV = 10 × 10 cm); In RAYSCAN α+, large jaw mode (field-of-view, FOV = 16 × 10 cm), jaw mode (FOV = 10 × 10 cm). The detailed exposure conditions for each mode were described in Table 1. The machine geometry for MC simulation were also described as suggested by individual manufacturer.

**Table 1.**
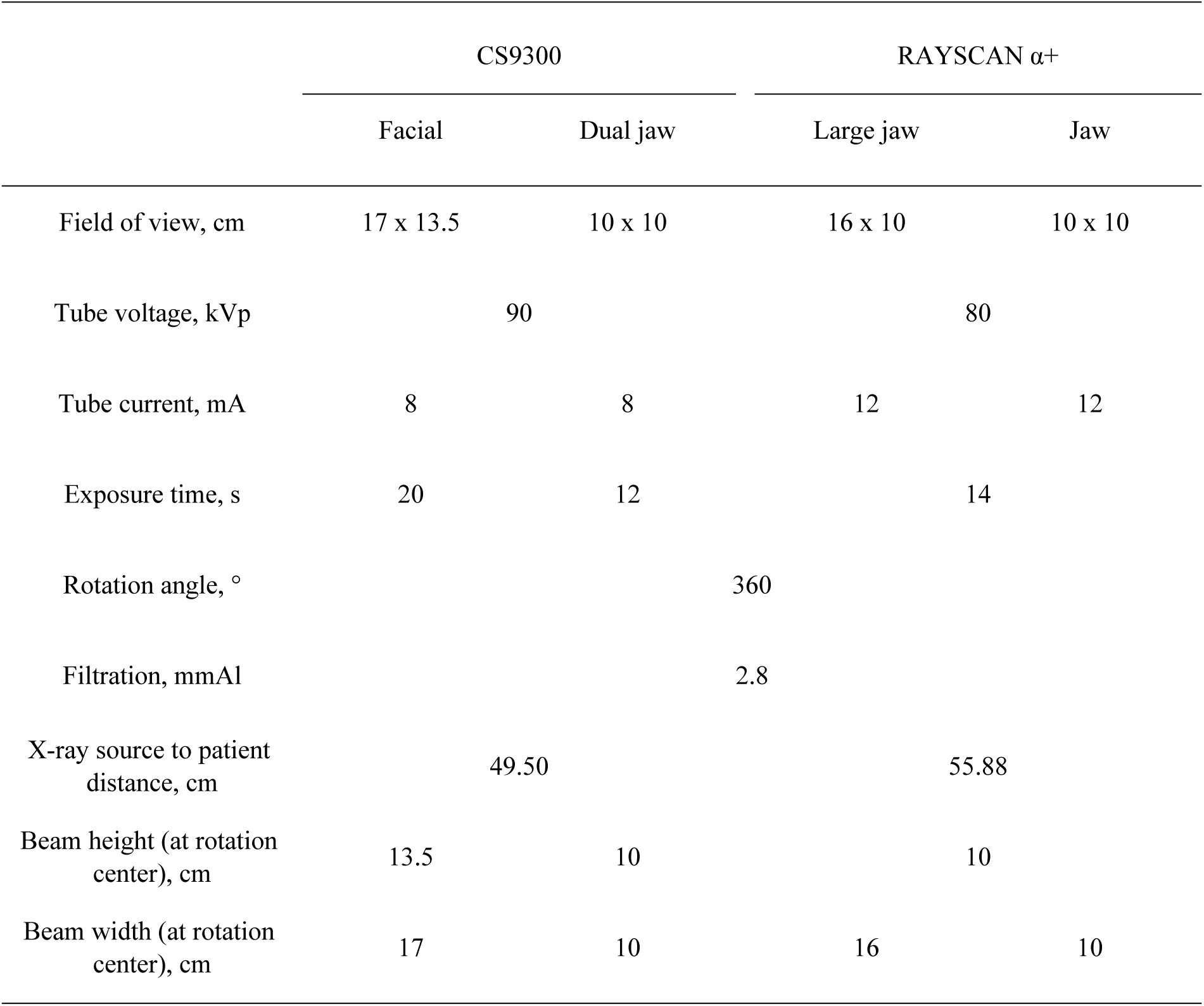
Exposure conditions of different modes in the CS9300 (Carestream Dental LLC, Atlanta, Georgia) and RAYSCAN α+ (Ray Co. Ltd, Hwaseong-si, Korea)

### 2. OSLD measurement

OSLD is a plastic disk containing aluminum oxide doped with carbon (Al2O3:C). This dosimetry absorbs radiation and this stored energy can be read out with light stimulation [12]. The dosimetry efficiently releases stored energy when stimulate with light of 540nm, still wide range of light can stimulate energy release thus the disk was encased in plastic holder. Each holder case was tagged with quick response (QR) code for identification of respective OSLD (Fig 1a).

**Figure 1.**
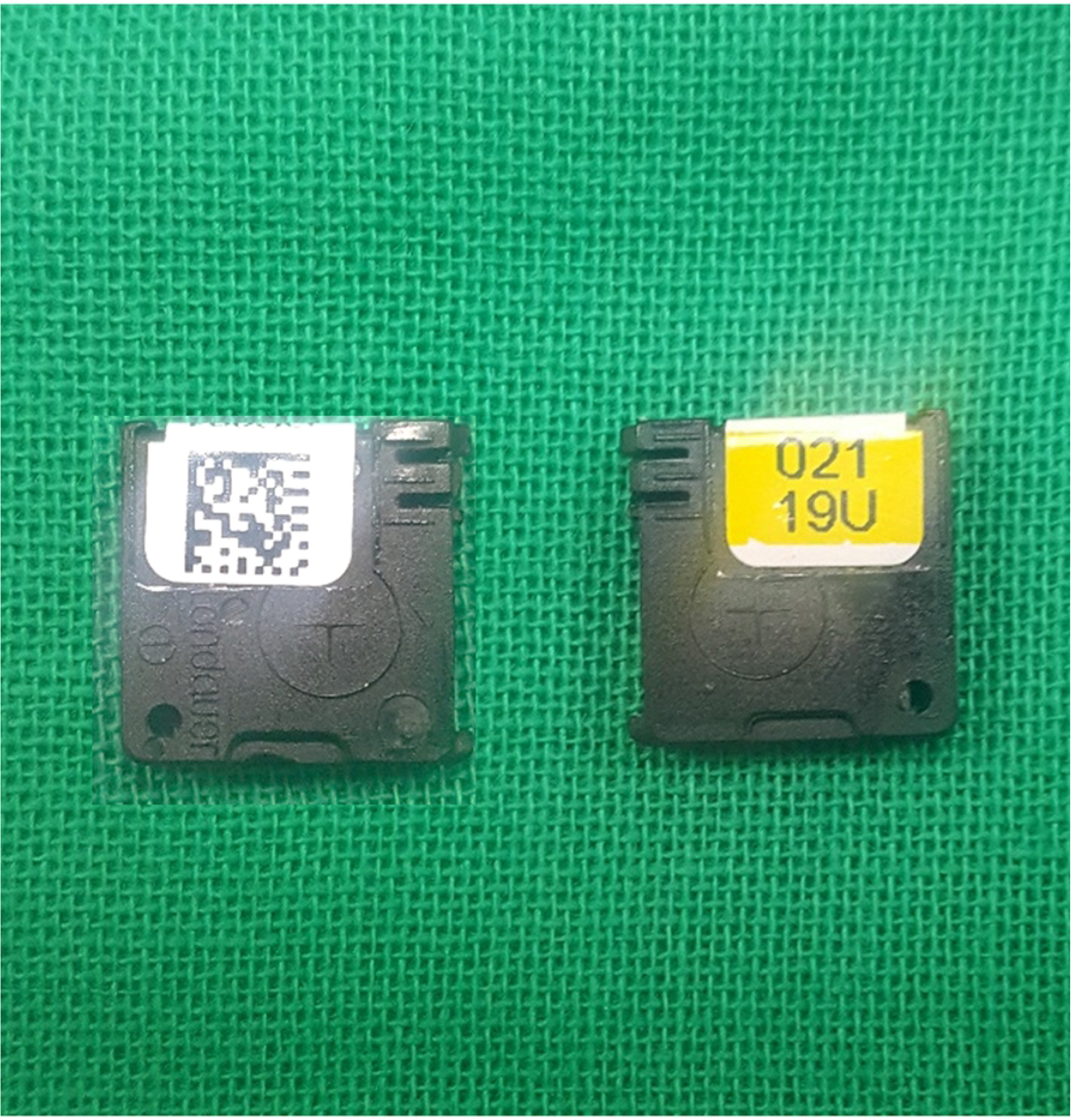
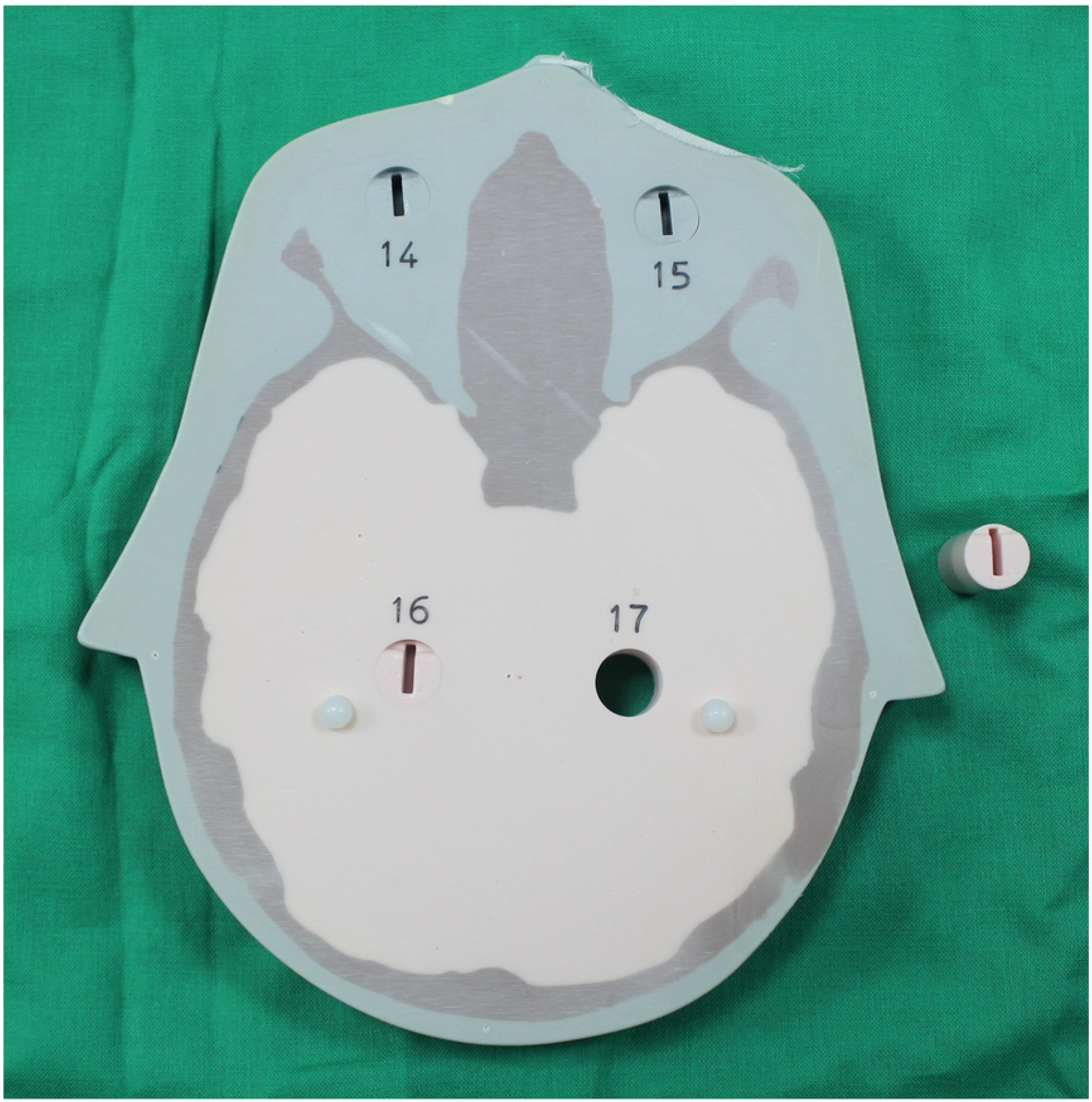
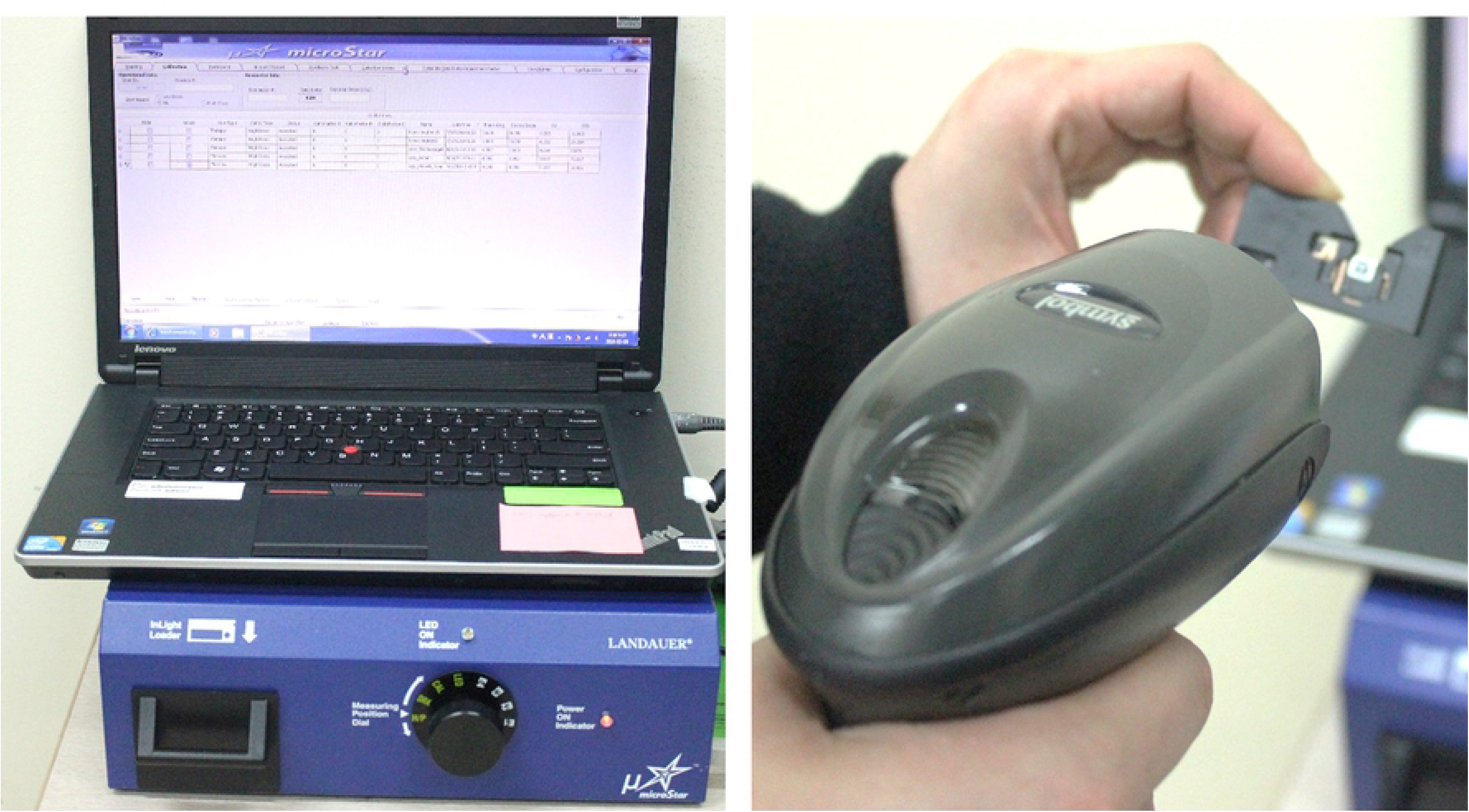
Experimental setting and facilitation for optically stimulated luminescence dosimeter (OSLD) measurement. (a) OSLD encased in a holder preventing light exposure. There is identification quick response (QR) code and identificaiton number marked on the case. (b) Human tissue equivalent phantom with dosimetry slot. (c) Dosimetry reader (MicroStar; Landauer) prepared optimal for 80 kilovoltage and low dose type (<30 mGy). Each dosimetry is identified with QR code and can be read out.

Total 22 OSLDs (nano-Dot, Landauer, Inc., Glenwood, IL) were placed in head and neck organs of adult head phantom (ATOM, CIRS, Norfolk, VA). This phantom was composed of tissue equivalent material and for each anatomy, there was slot for dosimetry placement (Fig. 1b). Details of the OSLD locations and corresponding tissues were described in Fig 2.

**Figure 2.**
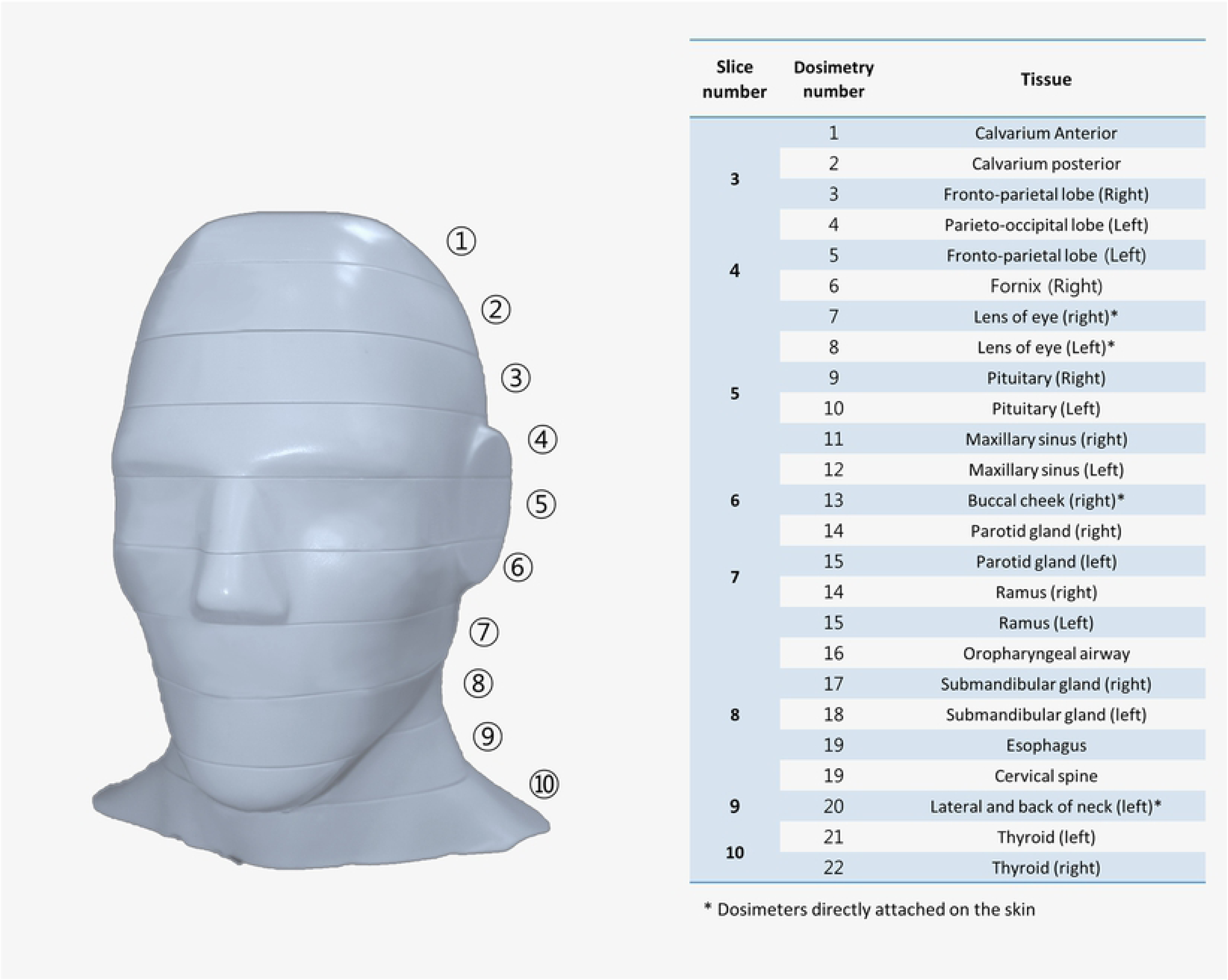
The location of optically stimulated luminescent dosimetry (OSLD) in an adult head and neck phantom (ATOM; CIRS, Norfolk, VA, USA) with the slice number of the phantom.

Phantom equipped with OSLD was exposed with four different examination modes (facial, dual jaw, large jaw and jaw) of two different units (CS9300 and RAYSCAN α+). All exposures were performed for twice and read dose values were averaged for the further calculation.

The reader (MicroStar; Landauer) was prepared to be optimized for 80 kilovoltage and low dose type (<30 mGy) and each dosimetry was identified with QR code and read out (Fig. 1c). The values were acquired as a photon counts with an accuracy of approximate ±2 % and this was converted to the dose in mGy unit using an energyspecific conversion factor. Those were converted into organ dose, mostly following the method done by Loudlow et al [8].

When multiple OSLDs were used for one organ, the average value was used. For example, the mean value of fronto-parietal lobe, parieto-occipital lobe, fornix and pituitary were used for brain dose. Bone marrow dose was obtained considering its distribution in mandible (0.8 %), calvaria (7.7 %) and cervical spine (3.8%) [13]. Bone surface dose were obtained with using coefficient, bone-to-muscle attenuation ratio, multiplied with bone marrow value. The equation for the coefficient was as followed: −0.0618 × kV(p) × 2/3 + 6.9406 [14]. The irradiated proportion of skin, lymphatic nodes and muscles on head and neck region are estimated as 5% and esophagus as 10 % of the whole body and this was taken consideration in organ dose calculation (Table 2) [15].

**Table 2.** Estimated fraction irradiated in tissues and tissue weighting factors recommended by the International Commission on Radiological Protection (ICRP)

|  | Fraction irradiated (%) | Tissue weighting factor | OSLD ID |
| --- | --- | --- | --- |
| Bone marrow | 12.2 | 0.12 |  |
| Mandible | 0.8 |  | 14, 15 |
| Calvaria | 7.7 |  | 1, 2 |
| Cervical spine | 3.8 |  | 19 |
| Thyroid | 100 | 0.04 | 21, 22 |
| Esophagus | 10 | 0.04 | 16 |
| Skin | 5 | 0.01 | 13, 20 |
| Bone surface* | 16.5 | 0.01 |  |
| Mandible | 1.3 |  | 14, 15 |
| Calvaria | 11.8 |  | 1, 2 |
| Cervical spine | 3.4 |  | 21 |
| Salivary glands | 100 | 0.01 |  |
| Parotid | 100 |  | 14, 15 |
| Submandibular | 100 |  | 17, 18 |
| Brain | 100 | 0.01 | 3, 4, 5, 6, 9, 10 |
| Remainder tissue |  | 0.12 |  |
| Lymphatic nodes | 5 |  | 14, 15, 17, 18, 19 |
| Muscle | 5 |  | 14, 15, 17, 18, 19 |
| Extrathoracic airways | 100 |  | 14, 15, 17, 18, 16 |
| Oral mucosa | 100 |  | 14, 15, 16, 17, 18, 19 |
| Eyes | 100 |  | 7, 8 |

\* Bone surface = bone marrow dose × bone/muscle mass energy absorption coefficient ratio (MEACR), MEACR = 0.0618 × 2/3 kVp + 6.9406 [14].

The organ doses were further integrated into the effective dose considering tissue weighted factor provided by International Commission on Radiological Protection (ICRP) 2007 (Table 2) [8, 16]. The equation for the effective dose calculation is as followed; E = Σ WT X HT, where E is effective dose, WT is the tissue weighting factor and HT is radiation weighed organ dose [16].

### 3. Monte Carlo simulation

Monte Carlo (MC) simulation is a widely used technique in the probabilistic analysis where random numbers are used for simulating the transport of radiation in complex medium such as human body [17]. When the physical information about x-ray examination technique was given, computer calculates organ absorbed dose with a MC simulation. In this study, commercial software commonly used in medical radiation dose calculation, PCXMC20Rotation (STUK, Helsinki, Finland) was used. According to the software manual, following factors were set for the software running; input dose, reference point, x-ray tube voltage, filtration, source-to-reference distance, x-ray beam width and height at reference point.

As an input dose, the exposure dose from the unit, dose-area-product (DAP, mGy·cm2) was selected and measured with DAP meter (VacuDAPTM; VacuTec Meßtechnik GmbH, Dresden, Germany). For respective examination mode, measurement was performed twice and the mean value was used (Fig 3). The reference point, the center of the x-ray unit during rotating through which all x-ray beams pass, was referenced to the previous literatures and marked as 3 dimensional coordination on X, Y, Z-axis (Fig. 4) [5, 10]. In addition, x-ray tube voltage, filtration and source-to-reference distance, beam width and height at reference point, were according to each examination mode in the specification of individual CBCT unit, provided by manufacturer (Table 1).

**Figure 3.**
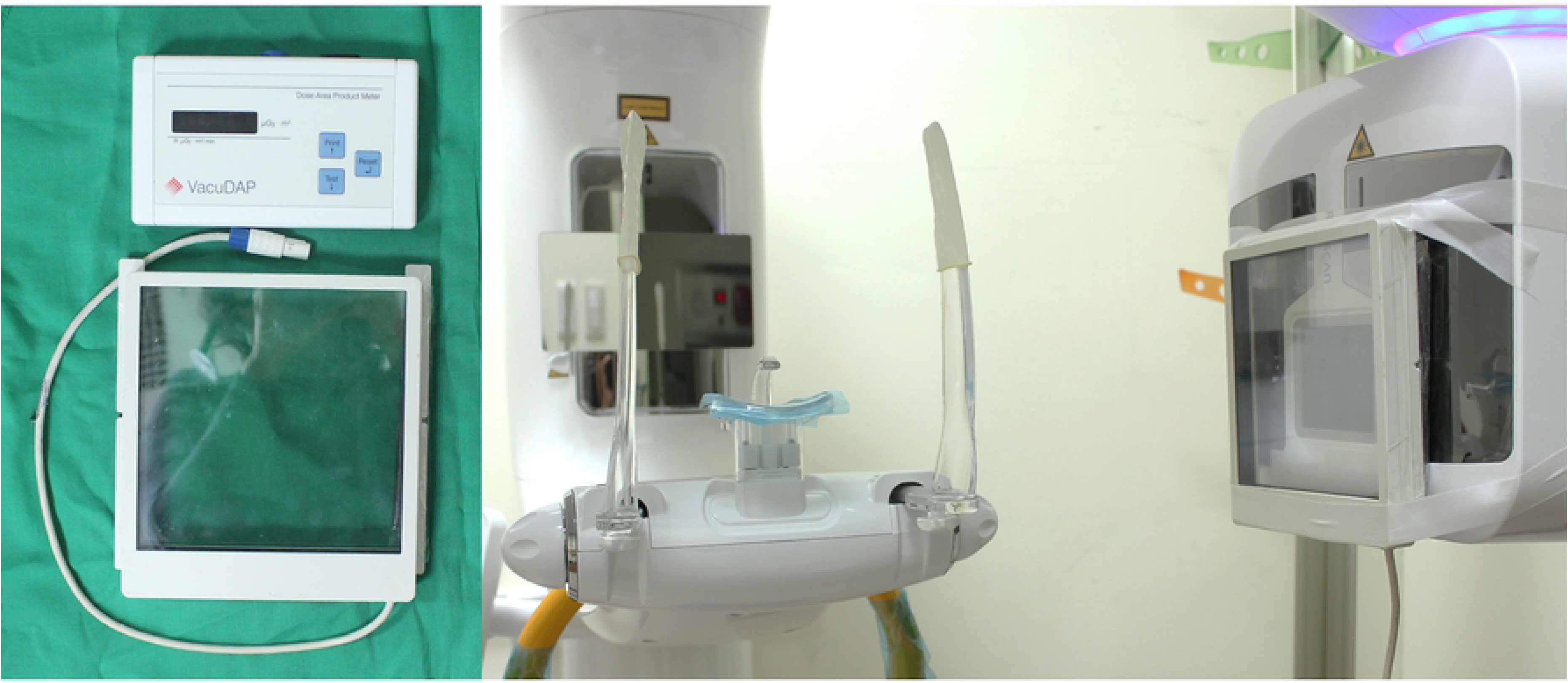
Dose-area-product (DAP) meter (VacuDAPTM; VacuTec Meßtechnik GmbH, Dresden, Germany) for input dose measurement. Ion chamber attached on the surface of x-ray tube head for the measurement.

**Figure 4.**
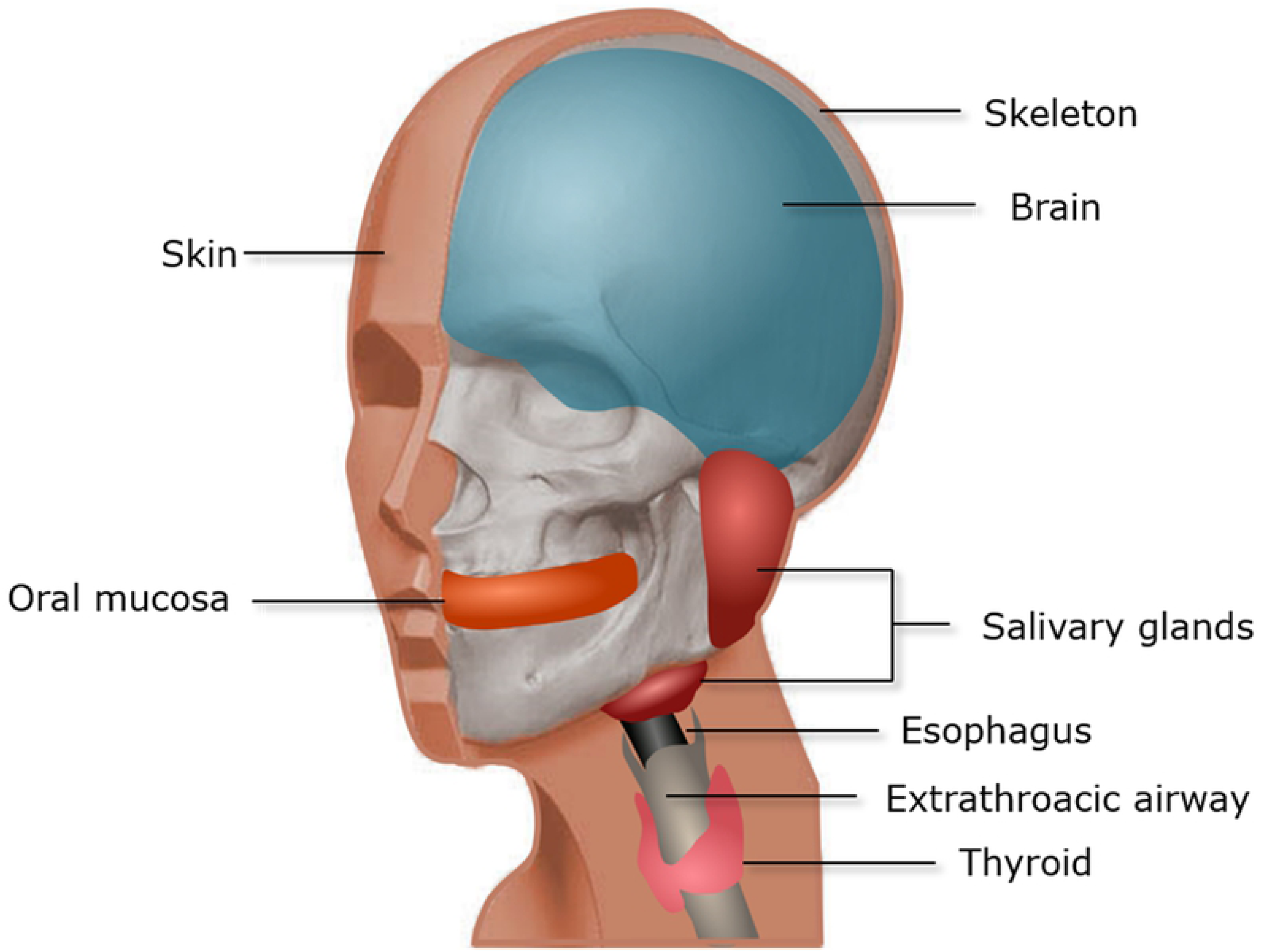
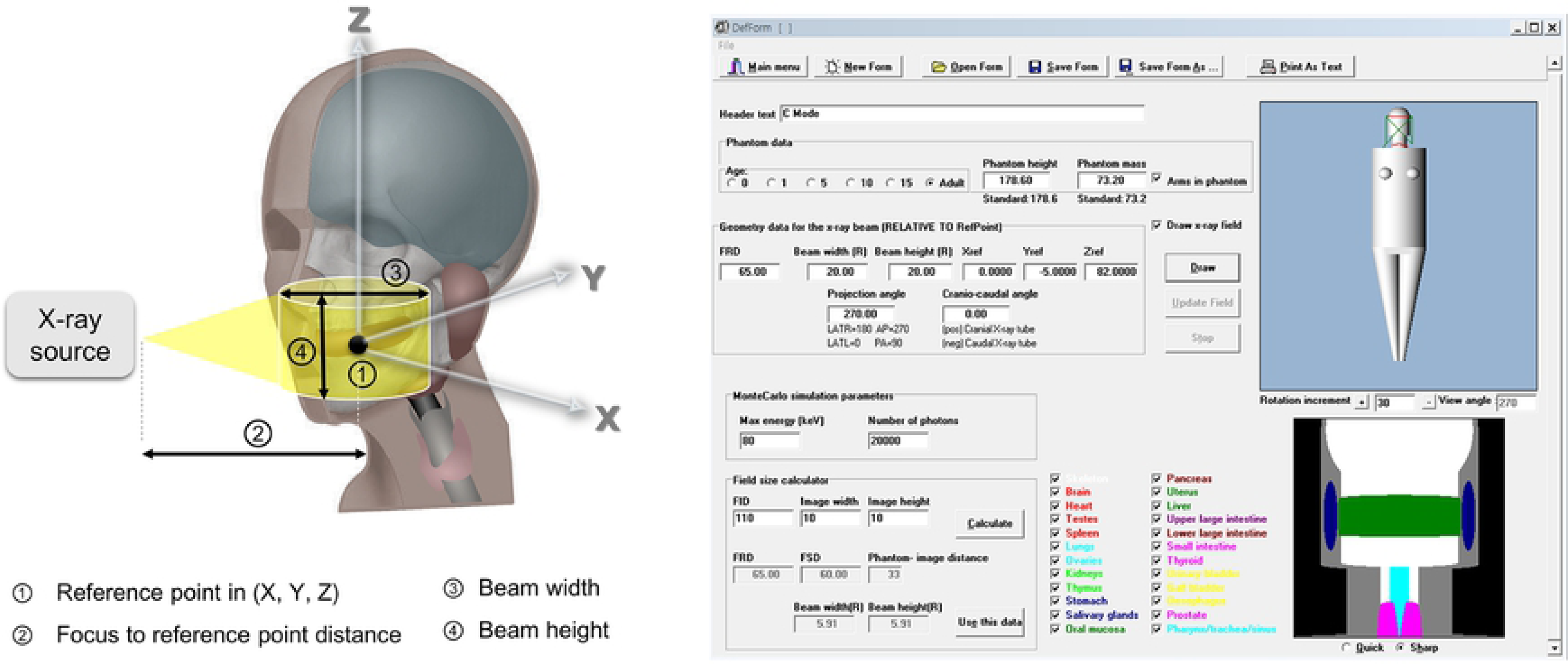
Virtual phantom and monte carlo simulation software. (a) Head and neck organs included in the virtual phantom. (b) Geometric variables required for the MC simulation and PCXMC20Rotation (STUK, Helsinki, Finland) software used in this study.

## Result

The mean DAP value measured with DAP meter was 215.1, 91.0, 176.6 and 167.9 mGy• cm^2^ respectively for facial, dual jaw, large jaw and jaw mode (Table 3). OSLD measured effective dose showed tendency of high value compared to that obtained with MC method. Only Dual jaw mode of CS9300 showed higher effective dose in MC method compared to OSLD method. The percent difference between the two methods was in the range of 4.0 to 14.3 %. The dose difference between the methods was decreased as the examination FOV decreased (Table 4). Organ dose were varied according to the method, while overall trend was similar in both methods (Fig 5). In other words, the organs irradiated relatively low dose in OSLD method mostly showed low dose in MC method. Organs with high dose in OSLD method also showed high dose in MC method. In both method, oral mucosa and salivary gland were two most irradiated organs (Fig 5).

**Table 3.** Mean and standard deviation of Dose-Area-Product (DAP) value measured with DAP meter (mGycm^2^) in different mode and units of cone-beam computed tomography

| CS9300 | | RAYSCAN $\alpha$+ | |
| --- | --- | --- | --- |
| Facial (17 x 13.5 cm) | Dual jaw (10 x 10 cm) | Large jaw (16 x 10 cm) | Jaw (10 x 10 cm) |
| 215.1 ± 0.4 | 91.0 ± 0.4 | 176.6 ± 0.4 | 167.9 ± 0.6 |

**Table 4.** The effective dose obtained with the OSLD and MC methods, and the percent difference.

| Effective dose (μSv) |  |  |  |  |
| --- | --- | --- | --- | --- |
|  | CS9300 |  | RAYSCAN α+ |  |
|  | Facial<br>(17 x 13.5 cm) | Dual jaw<br>(10 x 10 cm) | Large jaw<br>(16 x 10 cm) | Jaw<br>(10 x 10 cm) |
| OSLD method | 181.4 | 90.7 | 228.5 | 213.8 |
| MC method | 160.9 | 94.4 | 198.0 | 195.2 |
| Percent difference (%)* |  |  |  |  |
|  | CS9300 |  | RAYSCAN α+ |  |
|  | Facial<br>(17 x 13.5 cm) | Dual jaw<br>(10 x 10 cm) | Large jaw<br>(16 x 10 cm) | Jaw<br>(10 x 10 cm) |
|  | 12.0 | 4.0 | 14.3 | 9.1 |

\* Percent difference = $\left| \frac{effective\ dose\ (OSLD\ method) - effective\ dose\ (MC\ method)}{\frac{1}{2} \times effective\ dose\ (OSLD\ method) + effective\ dose\ (MC\ method)} \right| \times 100$

**Figure 5.**
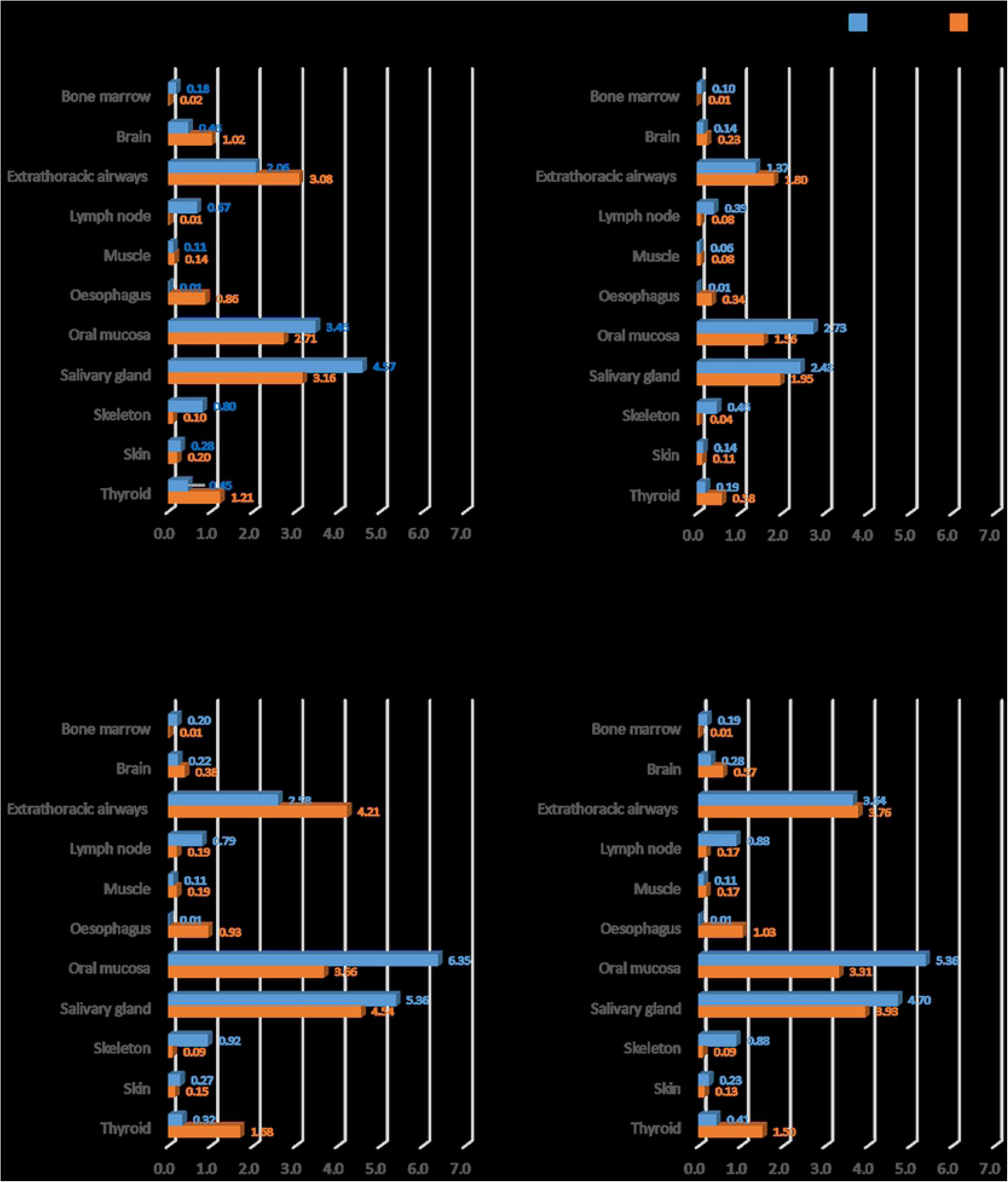
Organ dose of both methods according to the different CBCT unit and examination mode. Note that the values varied according to each organ, while the overall trend was similar in both methods.

## Discussion

Since the development of CBCT in the dental field, the usage of it has been growing rapidly, and research on its radiation dose has always been of interest. For now, various CBCT models from numerous manufacturer equipped with different exposure mode. In other words, exposure dose as well as patient absorbed or effective dose are varied on each CBCT machines of different examination mode. In 2015, Ludlow et al. meta-analyzed the effective dose study of CBCT conducted with dosimetry measurement. The value was widely varied from 46 to 1073 μSv for large field of view (FOV) and 9-560 μSv for medium FOV in each machines [3]. Assumed facial and large jaw mode as large and dual jaw and jaw mode as medium, our study, both OSLD and MC method, showed effective dose included in this range, regardless of the method. The major contribution to the wide range of effective dose in different CBCT units of similar FOV would be probably different exposure conditions of each machines, however, different dose measurement method also influenced to raise deviations of overall effective dose assessment [3]. Thus, consensus in dose evaluation method is consequently needed for comparative analysis in effective dose reporting of each machine. This consequently helps to construct database of patient dose and setting nation-wide regulation for the CBCT dose. Ludlow et al. studied the effective dose with OSLD and the same CBCT unit used in this study, CS9300, and reported 204 and 76 μSv respectively for facial and dual jaw mode [3]. Even though the same method and materials were used for dose evaluation, the effective dose values showed differences even greater or similar than that between OSLD and MC method. This was probably caused by the sampling error, as it was also mentioned by Loudlow et al [3]. The sampling error is defined as the influence of location, distribution and the number of dosimeter used in each organ to the measured value. It is difficult to use the same number of dosimetry in every experiment performed by different experimenters, due to the practical reason such as cost of dosimetry. Also, phantom positioning within the CBCT unit during the exposure is another challenging part causing large deviation in resulting organ dose and the effective dose.

On such aspect, application of MC method might be more reproducible and practical while accurate. First of all, the effective dose obtained with MC simulation was relatively good agreement with that obtained with OSLD that the percent differences were under 15 %. Toivonen et al.[18] assumed as ‘good’ in agreement when the difference between the dosimetry and computer simulated methods is below 25 %. Second, user dependent factors were limited in MC method during the whole measurement process. According to the previous studies adopted MC method for dose evaluation, machine geometry, such as filtration, tube voltage, x-ray beam width or height and source-to-subject distance, are the information required for the simulation [10, 19, 20]. For the current study, manufacturer of the machine provided required information in the specification. This method is also efficient in that it cost less than preparing human tissue-equivalent phantom, dosimetry, and dosimetry reading device.

There is important consideration for adopting MC method currently. The virtual phantom used for the simulation should be standardized [21]. In 2009, ICRP introduced reference phantom of female and male adult which is based on the actual computed tomographic data of adult human [22]. Among previous studies, only one adopted ICRP reference phantom and others used computed tomographic scan data of Rando-alderson phantom [11, 23]. In present study, the Cristy and Eckerman phantom facilitated in software was used without any modification. The Cristy and Eckerman phantom was describing human body organs as simplified form using cone, ball or cylinder shape. Compared to the ICRP reference phantom, it is not sophisticated enough to simulate precise organ absorbed and effective dose in dental CBCT, exposing relatively low dose compared to medical CT.

This probably contributed to the difference between the dose from OSLD method and MC simulation, as well, in this study. Ludlow et al.[3] in 2015 mentioned that DAP is not appropriate to be used for obtaining effective dose. This statement is true, if we simply convert DAP value into effective dose using converting coefficient. Several studies attempted to find converting coefficient to obtain effective dose of CBCT, while coefficients are different by individual CBCT model with unique geometry [24, 25]. On the contrary, MC simulation calculates the effective dose taking individual machine geometry into the consideration. Thus, it may produce more precise results based on the DAP value, compared to the simple conversion method of DAP value into effective dose.

In the present study, DAP measurement was performed using DAP meter. DAP measurement procedure is not experimenter specific, still, it requires equipment composed of ion chamber, DAP meter and cables. Also, the procedure takes time and experimenter’s labor. Fortunately, recent CBCT machines provide DAP value according to the exposure condition. Though, this value is not real-time measured, and predetermined value by the manufacturer, MC simulation software with precise reference phantom, it may be possible to obtain an approximate effective dose which is not depended on the experimenter or the measurement method.

In conclusion, the effective dose by individual CBCT models and examination modes is continuously reported and large data has been accumulated up to now [3, 26]. To contribute for this big data accumulation, the effective dose obtained by two different methods and CBCT machines was reported in this study. The ultimate goal of the effective dose assessment and data accumulation is a dose reduction and regulation for patient’s benefit. To attain this, more importantly, consensus in dose evaluation method is essential. In addition, development of a relatively accurate and easy-handling method would contribute more dose data acquisition. Therefore, we carefully suggest MC simulation based on reference phantom for further dose evaluation.

## Acknowledgement

This research was supported by Basic Science Research Program through the National Research Foundation of Korea (NRF) funded by the Ministry of Education (No. 2017R1D1A1B03033218)

## References

1. Hart D, Hillier M, Wall B. National reference doses for common radiographic, fluoroscopic and dental Xray examinations in the UK. The British journal of radiology. Br J Radiol. 2009;82(973):1–12.

2. Loubele M, Bogaerts R, Van Dijck E, et al. Comparison between effective radiation dose of CBCT and MSCT scanners for dentomaxillofacial applications. European journal of radiology. Eur J Radiol. 2009;71(3):461–8.

3. Ludlow J, Timothy R, Walker C, et al. Effective dose of dental CBCT—a meta analysis of published data and additional data for nine CBCT units. Dentomaxillofac Radiol. 2015;44(1):20140197

4. Yepes JF, Booe MR, Sanders BJ, et al. Pediatric Phantom Dosimetry of Kodak 9000 Cone-beam Computed Tomography. Pediatr Dent. 2017;39(3):229–232

5. Koivisto J, Kiljunen T, Tapiovaara M, Wolff J, Kortesniemi M. Assessment of radiation exposure in dental cone-beam computerized tomography with the use of metal-oxide semiconductor field-effect transistor (MOSFET) dosimeters and Monte Carlo simulations. Oral Surg Oral Med Oral Pathol Oral Radiol. 2012;114(3):393–400.

6. Sarrabayrouse G, Siskos S. Radiation dose measurment using MOSFETs. IEEE Instrumentation & Measurement Magazine. 1998;1(2):26–34.

7. Jursinic PA. Characterization of optically stimulated luminescent dosimeters, OSLDs, for clinical dosimetric measurements. Med Phys. 2007;34(12):4594–604.

8. Ludlow JB, Walker C. Assessment of phantom dosimetry and image quality of i-CAT FLX conebeam computed tomography. Am J Orthod Dentofacial Orthop. 2013;144(6):802–17.

9. Morant JJ, Salvadó M, Casanovas R, Hernández-Girón I, Velasco E, Calzado A. Validation of a Monte Carlo simulation for dose assessment in dental cone beam CT examinations. Phys Med. 2012;28(3):200–9.

10. Lee C, Lee S-S, Kim J-E, et al. Comparison of dosimetry methods for panoramic radiography: thermoluminescent dosimeter measurement versus personal computer–based Monte Carlo method calculation. Oral Surg Oral Med Oral Pathol Oral Radiol. 2016;121(3):322–9.

11. Ernst M, Manser P, Dula K, Volken W, Stampanoni M, Fix M. TLD measurements and Monte Carlo calculations of head and neck organ and effective doses for Cone Beam Computed Tomography using 3D Accuitomo 170. Dentomaxillofac Radiol. 2017;46(7):20170047.

12. Jursinic PA. Changes in optically stimulated luminescent dosimeter (OSLD) dosimetric characteristics with ccumulated dose. Med Phys. 2010;37(1):132–40.

13. Cristy M. Active bone marrow distribution as a function of age in humans. Phys Med Biol. 1981;26(3):389–400.

14. International Commission on Radiation Units and Measurements. Physical Aspect of Radiation. Washington, DC 1963.

15. Ludlow J, Davies-Ludlow L, Brooks S. Dosimetry of two extraoral direct digital imaging devices: NewTom cone beam CT and Orthophos Plus DS panoramic unit. Dentomaxillofac Radiol. 2003;32(4):229–34.

16. Valentin J. The 2007 recommendations of the international commission on radiological protection: Elsevier Oxford; 2007.

17. Tapiovaara M, Siiskonen T. PCXMC, A Monte Carlo program for calculating patient doses in medical x-ray examinations. J ICRU. 2005;5(2):100–2.

18. Toivonen M, Aschan C, Rannikko S, Karuka K, Savolainen S. Organ dose determinations of x-ray examinations using TL detectors for verification of computed doses. Radiat Prot Dosimetry. 1996;66(1-4): 298–294.

19. DeMarco J, Cagnon C, Cody D, et al. A Monte Carlo based method to estimate radiation dose from multidetector CT (MDCT): cylindrical and anthropomorphic phantoms. Phys Med Biol. 2005;50(17):3989–4004.

20. Podnieks E, Negus I. Practical patient dosimetry for partial rotation cone beam CT. The British journal of radiology. Br J Radiol. 2012;85(1010):161–7.

21. Zhang G, Marshall N, Bogaerts R, Jacobs R, Bosmans H. Monte Carlo modeling for dose assessment in cone beam CT for oral and maxillofacial applications. Med Phys. 2013;40(7):072103

22. ICRP. Adult Reference Computational Phantoms. Vol ICRP Publication 1102009.

23. Morant J, Salvadó M, Hernández-Girón I, Casanovas R, Ortega R, Calzado A. Dosimetry of a cone beam CT device for oral and maxillofacial radiology using Monte Carlo techniques and ICRP adult reference computational phantoms. Dentomaxillofac Radiol. 2013;42(3):92555893.

24. Kim D-S, Rashsuren O, Kim E-K. Conversion coefficients for the estimation of effective dose in cone-beam CT. Imaging science in dentistry. Imaging Sci Dent. 2014;44(1):21–9.

25. Shin H, Nam K, Park H, Choi H, Kim H, Park C. Effective doses from panoramic radiography and CBCT (cone beam CT) using dose area product (DAP) in dentistry. Dentomaxillofac Radiol. 2014;43(5):20130439.

26. Pauwels R, Beinsberger J, Collaert B, et al. Effective dose range for dental cone beam computed tomography scanners. European journal of radiology. Eur J Radiol. 2012;81(2):267–71.

